# An optimal population code for global motion estimation in local direction-selective cells

**DOI:** 10.1101/2021.03.17.435642

**Authors:** Miriam Henning, Giordano Ramos-Traslosheros, Burak Gür, Marion Silies

**Affiliations:** Institute of Developmental Biology and Neurobiology, Johannes-Gutenberg University Mainz, 55128 Mainz, Germany; Göttingen Graduate School for Neurosciences, Biophysics, and Molecular Biosciences (GGNB) and International Max Planck Research School (IMPRS) for Neurosciences at the University of Göttingen, 37077 Göttingen, Germany

**Keywords:** global motion processing, convergent evolution, population code, neuroethology, efficient coding, direction-selectivity, Drosophila

## Abstract

Nervous systems allocate computational resources to match stimulus statistics. However, the physical information that needs to be processed depends on the animal’s own behavior. For example, visual motion patterns induced by self-motion provide essential information for navigation. How behavioral constraints affect neural processing is not known. Here we show that, at the population level, local direction-selective T4/T5 neurons in *Drosophila* represent optic flow fields generated by self-motion, reminiscent to a population code in retinal ganglion cells in vertebrates. Whereas in vertebrates four different cell types encode different optic flow fields, the four uniformly tuned T4/T5 subtypes described previously represent a local snapshot. As a population, six T4/T5 subtypes encode different axes of self-motion. This representation might serve to efficiently encode more complex flow fields generated during flight. Thus, a population code for optic flow appears to be a general coding principle of visual systems, but matching the animal’s individual ethological constraints.

## Introduction

Evolution has matched neural resources to the sensory information that is available to the animal (1, 2). This is particularly well studied in vision, where the sensitivity of photoreceptors efficiently covers the range of intensities in the environment (3, 4), and many other specializations in retinal circuitry match the statistics of visual information (5, 6). But how did evolution accommodate sensory systems in which the physical distribution of information encountered depends on animal behavior? For example, any visual animal that navigates through its environment needs to detect and compute global motion patterns elicited on the eye, which will depend on the type of locomotion. In vertebrates, such optic flow generated by self-motion is represented by the population of local motion-sensitive retinal ganglion cells(7). Here, topographically organized directional tuning maps represent optic flow that match the global motion patterns generated during walking. In insects, the first direction-selective cells that encode local motion are the T4 and T5 neurons of the ON and OFF pathways. T4/T5 are thought to be uniformly tuned throughout the visual field, representing the four cardinal directions: upward, downward, front-to-back and back-to-front motion(8–10). One synapse downstream of T4/T5 cells, optic flow patterns are then encoded by wide-field neurons that sample information globally across visual space(11, 12). This suggests that the coding of optic flow is fundamentally different between vertebrate and invertebrate visual systems. It is unclear why flies would have evolved a system in which optic flow has to be computed through complex transformations from local motion detectors with uniform tuning, to ultimately match the motion patterns generated during flight.

Insect wide-field neurons that are tuned to specific optic flow patterns generated by forward movement or turns of the animal have been extensively characterized(13–15). Such wide field neurons, the lobula-plate tangential cells (LPTCs), also exist in *Drosophila*(16–18), and are involved in the control of optomotor responses, as well as in stabilizing gaze and forward walking(19–21). To extract optic-flow information, wide-field neurons pool information from presynaptic local motion detectors. In *Drosophila*, this is achieved by LPTCs receiving strong input from the columnar T4/T5 direction-selective cells (8, 22). T4/T5 provide excitatory input to downstream LPTCs within the same layer and inhibitory input via lobula plate-intrinsic neurons to LPTCs of the adjacent lobula plate layer with opposite tuning, thus establishing motion opponency (22, 23). Most LPTCs extend their dendrites along one layer of the lobula plate and thus pool information from one subtype of T4/T5 neurons (16, 17, 24), although recently described LPTCs also project to more than one layer (18, 25). Additionally, local motion signals are selectively amplified within the LPTC dendrites if they match the preferred global motion pattern (26). However, it is not fully understood how the flow-field-encoding receptive fields in the LPTCs are computed from presynaptic circuitry. The direction-selective T4/T5 cells respond to local motion, together forming a retinotopic map (8, 9). Whether their tuning to four cardinal directions generalizes over retinotopic locations is not known.

Here, we characterize the direction tuning distribution of T4/T5 neurons across anatomical and visual space. We demonstrate that directional preference of T4/T5 subtypes changes gradually, forming continuous maps of tuning. At the population level, T4/T5 cells in fact do not fall into four but six subgroups that encode six diagonal directions of motion, matching the hexagonal lattice of the eye. The six topographic tuning maps match optic flows field generated by self-motion of the fly. Therefore, the organization of local direction-selective cells that represents self-motion parallels the retinal code for optic flow, providing a striking example of convergent evolution. The specific types of optic flow that are encoded differ between the mouse retina and the *Drosophila* visual system, arguing that evolution matched neural resources to the different physical distribution of information encountered during walking or flight.

## Results

### T4/T5 population tuning clusters around hexagonal directions of motion

To understand how the T4/T5 neurons contribute to downstream optic flow fields, it is necessary to have a detailed map of T4/T5 direction tuning across retinotopic space. We used *in vivo* two-photon calcium imaging to record motion responses from large populations of T4/T5 neurons in individual flies. We imaged GCaMP6f responses to ON and OFF edges moving in eight directions at different fly orientations relative to the screen, together subtending ∼150° in azimuth and ∼60° in elevation (**fig. S1, A and B**). Tuning across 3537 individual cells (1376 T4, 2161 T5), recorded in 14 flies was broad, together spanning 360° of motion. Neurons in both layer A and B covered more than 120° of tuning direction, and thus twice the range of cells in layers C and D, which were tuned to a range of ∼60° (**Fig. 1A**). Dorsoventral location strongly impacted tuning direction in layers A and B (**Fig. 1A**). In layer A, cells that were more dorsally located in the lobula plate preferentially covered the 300°-360° range, whereas more ventral cells of the lobula plate showed tuning directions in the 0°-60° range. In layer B, more dorsally located cells were tuned to the 120°-180° range, and more ventrally located cells were tuned to 180°-240° (**Fig. 1A**). Although the population of T4/T5 cells covered all directions of motion, the tuning distribution was non-uniform (*Circular Rayleigh test*: *p<0*.*0001*).

**Fig 1.**
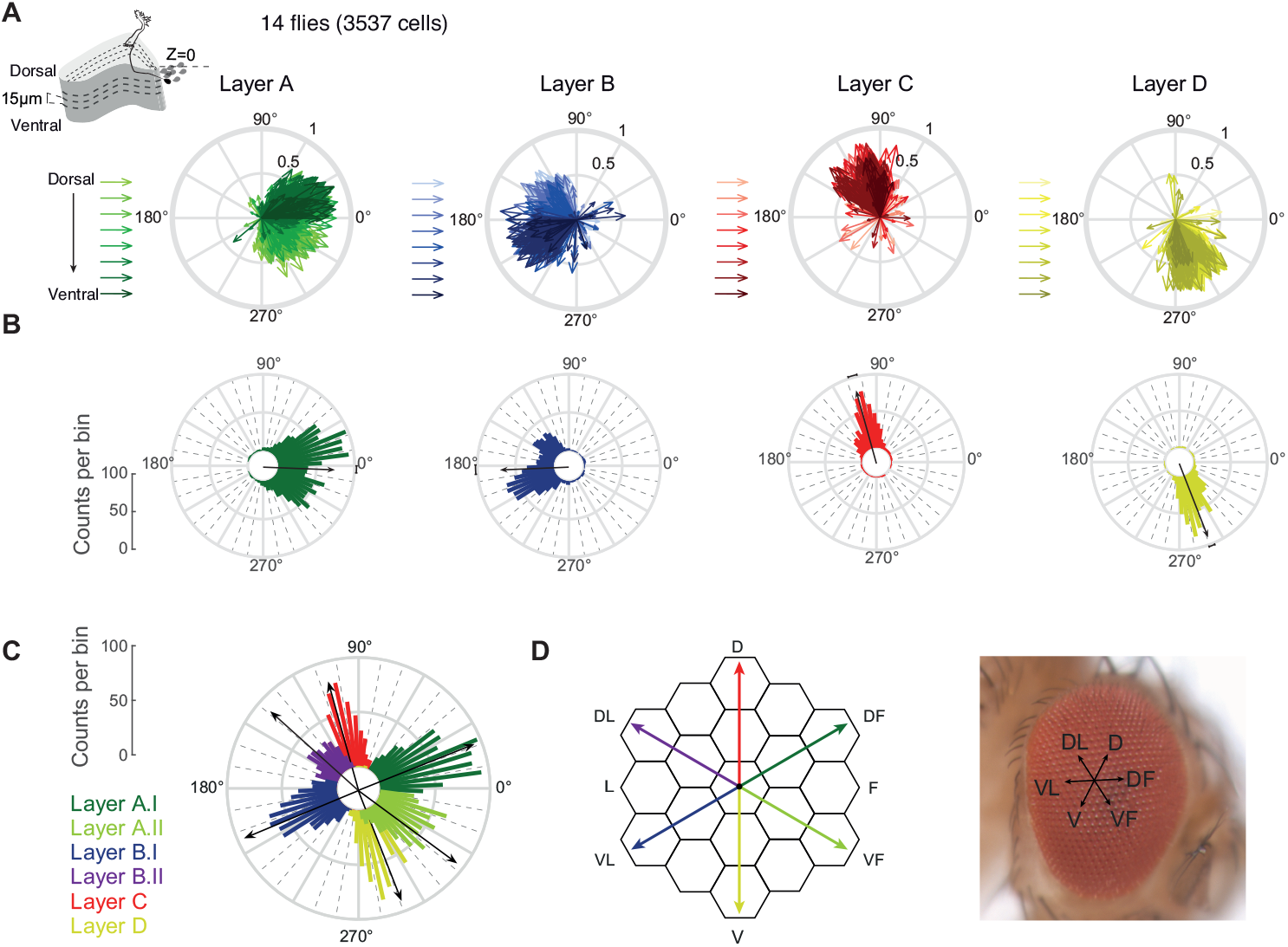
Directional tuning clusters around hexagonal directions of motion. (**A**) Directional tuning of individual neurons from 3537 cells (Layer A: 479/926 T4/T5. Layer B: 252/662 T4/T5. Layer C: 365/220 T4/T5. Layer D: 280/353 T4/T5) in 14 flies. Motion responses were represented by a vector(27), whose direction depicts tuning, whereas its length indicates selectivity. Hue illustrates z-depth relative to a reference (the outermost T4/T5 cell bodies). (**B** and **C**) Circular histograms of neuronal tuning preference. Black vectors depict average tuning per layer (B) or subtype (C). (**D**) On average, tuning of the six subtypes matches the hexagonal arrangement of the fly eye.

Looking at the number of neurons sensitive to a certain motion direction, most neurons in layers A and B were tuned to the diagonal directions of motion, flanking the overall average orthogonal tuning of these layers (**Fig. 1B**). Cells in layers C and D each showed a unimodal directional tuning distribution in the upward or downward direction, respectively (**Fig. 1B**). The bimodal distribution in layers A and B were well fit by two Gaussians (**fig. S1C**). When thus assigning each cell to one of six subtypes, tuning of two subtypes in layer A and B split at 0° or 180°, respectively (**fig. S1D**). The population average of the A.I subtypes was tuned to diagonal upward motion (∼30°) and the A.II subtype was tuned to diagonal downward motion (∼330°). Layer B subtypes encoded the two opposite axes of motion direction (**fig. S1D**). Taken together, our data show that at the population level, T4/T5 neurons fall into six functional subtypes (**Fig. 1C** and **fig. S1D**) Average motion tuning within individual subtypes reveals sensitivity to six directions with each subtype spanning a 60°-range, matching the hexagonal arrangement of the fly compound eye (**Fig. 1, C and D**).

To understand the spatial organization of six T4/T5 subtypes projecting to four anatomically distinguishable lobula plate layers, we plotted cellular subtype identity back onto the anatomical structure of the lobula plate (**Fig. 2A** and **fig. S2**). T4/T5 cells of one subtype dominated one lobula plate layer recorded in one plane along the dorsoventral axis. At more ventral planes, subtype A.I and B.I as well as the single respective subtypes of layers C and D were found more frequently. Dorsal planes more prominently housed subtypes A.II and B.II, but hardly showed any layer C or D cell responses (**Fig. 2, A and B** and **fig. S2**). This argues for a spatial separation of layer A and B subtypes at the level of T4/T5 axon terminals. Importantly, local T4/T5 recordings in an individual fly preferentially showed either four subtypes (as e.g. described in(8, 9)), or two subtypes, each representing snapshots of the T4/T5 population (**Fig. 2, A to D**). Only a global analysis of tuning revealed the six T4/T5 subtypes encoding six diagonal directions of motion (**Fig. 2e**).

**Fig 2.**
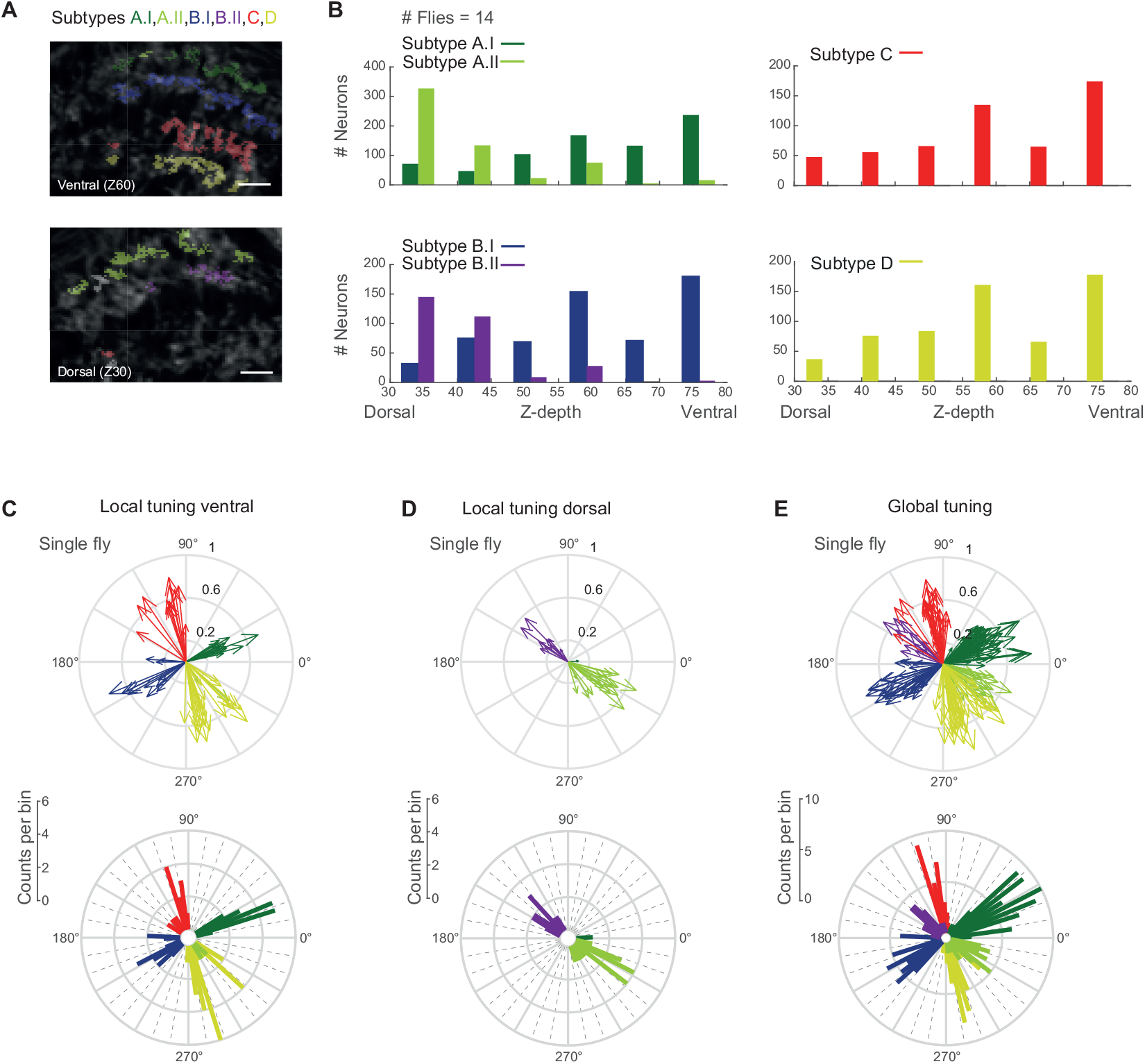
Layer A and B subtype projections separate along the dorsoventral axis. (**A**) *In vivo* two photon calcium images of the lobula plate at two planes along the dorsoventral axis (Z30/Z60 = z-depth 30/60 μm). ROIs are color coded based on their subtype identity. (**B**) Histograms displaying number of neurons from the different classes along the dorsoventral axis (z-depth). Scale bar 10 µm. (**C** and **D**) Tuning of individual neurons from one fly recorded in a ventral (Z60) (C), or dorsal (Z30) (D) plane of the lobula plate. Below: Same data plotted as circular histograms. (**E**) Tuning of all neurons recorded at different dorsoventral planes within one fly.

### T4/T5 neurons form topographic maps of directional tuning

We next asked if the ∼60° distribution of directional tuning within one subtype was random, or topographically organized. Color coding axon terminals based on their directional preference revealed that the tuning of neighboring cells was similar and gradually changed along the distal-to-proximal axis. As such, recording in one ventral plane of layer A (group A.I) revealed T4/T5 tuning ranging from diagonally upward on the proximal end to front-to-back motion on the distal end of the lobula plate (**Fig. 3, A and B**). T4/T5 cells of other subtypes also gradually changed tuning from proximal to distal. Subtler changes in the tuning of neighboring cells within one subtype were also apparent along the dorsoventral axis (**fig. S3, A and B**). This gradually distributed tuning existed for both T4 and T5 when analyzed separately (**Fig. 3, C and D** and **fig. S3C**). Because T4/T5 neurons are retinotopically organized, this directional tuning map suggests that the population of T4/T5 cells is sensitive to specific global motion patterns.

**Fig 3.**
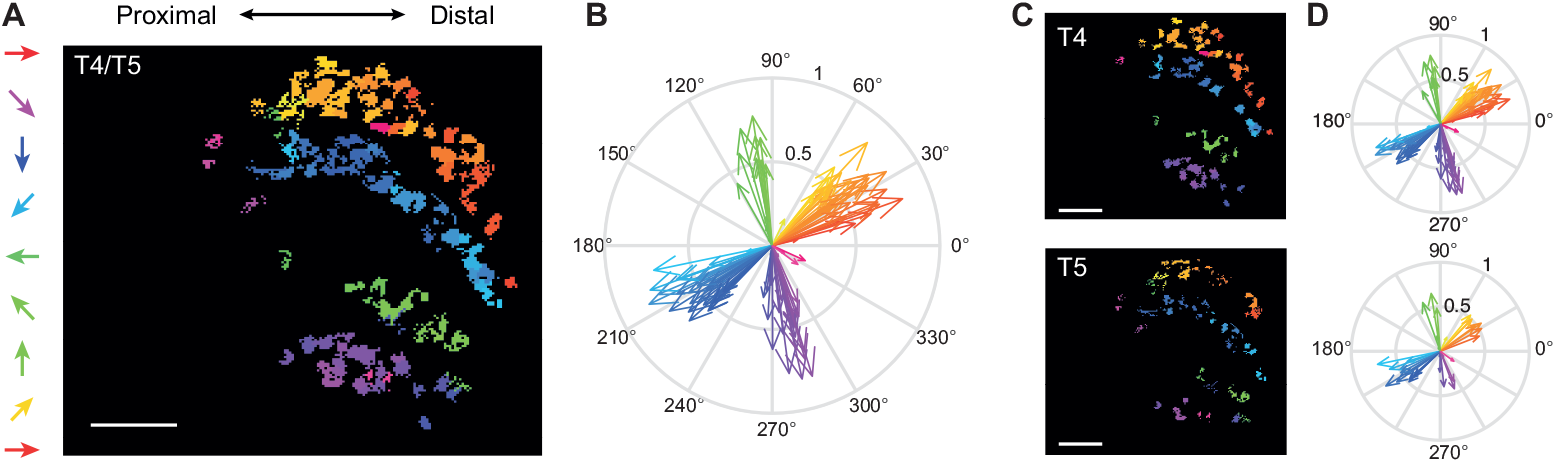
T4/T5 neurons form topographic maps of directional tuning. (**A**) Image of one layer of the lobula plate with ROIs color coded according to their directional tuning. (**B**) Data from (A) shown in vector space. (**C** and **D**) Data shown separately for T4 and T5 neurons. Scale bar = 10 µm.

### The six T4/T5 subtypes encode optic flow induced by self-motion

The local differences in the tuning preference within one T4/T5 subtype are reminiscent of direction-selective ganglion cells in the vertebrate retina, where the population of cells encodes translational optic flow generated by self-motion of the animal(7). We hypothesized that the differential tuning measured within each subtype of T4/T5 cells in the fly visual system serves a similar function. To relate tuning to the visual input, we mapped receptive-field centers (**fig. S4, A and B**) and plotted tuning at each receptive-field location on the screen (**fig. S4, C and D**). This revealed that cells of one subtype do not encode a uniform direction of motion, but rather that direction tuning of all cells within one subtype change gradually across visual space (**Fig. 4A** and **fig. S4E**). These topographic tuning maps resemble flow fields generated by different directions of self-motion in the fly. T4/T5 neurons in layers C and D appear to encode optic flow generated by downward or upward movement of the fly, whereas T4/T5 neurons in layers A and B seem to be tuned to diagonally upward or downward motion. The two flow fields encoded by the two subtypes of layers A or B are vertically flipped versions of each other (**Fig. 4A**). The successive change of tuning along azimuth and elevation matches the change of tuning seen in the topographic maps in the lobula plate (**Fig. 4A, Fig. 3**, and **fig. S4, E and F**).

**Fig 4.**
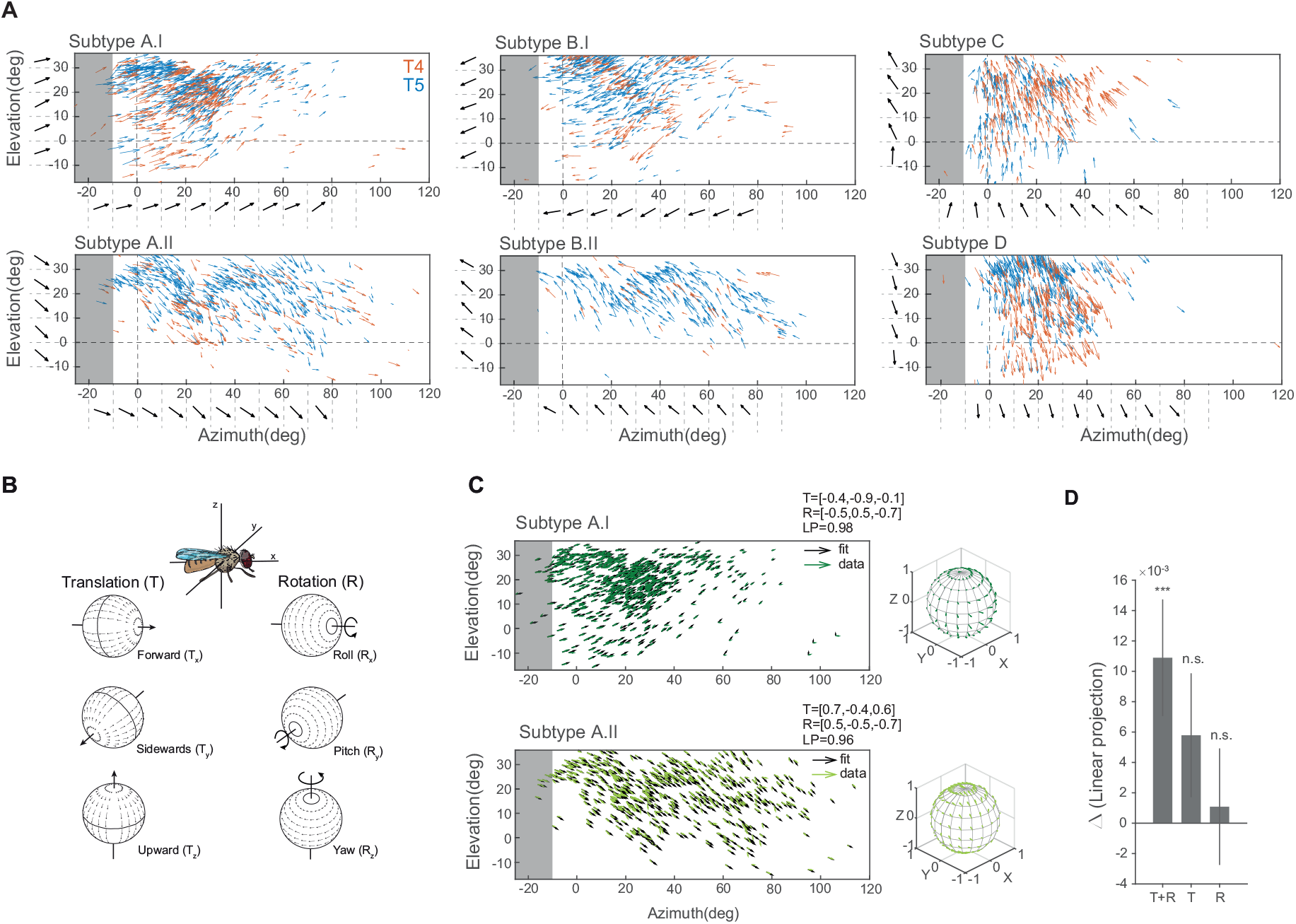
The population of T4/T5 neurons encode optic flow induced by self-motion. (**A**) Arrows indicate tuning direction of individual neurons plotted at their receptive field center coordinates in visual space. Length of vectors indicates direction selectivity of T4 (red) or T5 neurons (blue), n=14 flies. Horizontal and vertical dashed lines mark the split between left and right visual hemispheres and the horizon, respectively. Black tuning vectors show mean across 10 degree-wide bins. Gray shaded areas indicate the visual space in the left hemisphere that cannot be seen by the right eye. (**B**) Schematic of rotational and translational flow fields around the three body axes of the fly (modified after (29)). (**C**) Flow fields of data from two subtypes and the fitted normalized optic-flow model. Model vectors are shown for each corresponding data vector. (**D**) Differences of the fit quality for three model types (translation + rotation [T+R], pure translation [T], pure rotation [R]) compared to a uniform vector field model (***Wilcoxon: p*<*0.001).

To investigate the type of self-motion encoded by the different subtypes, we trained an optic flow model (28) to match the population receptive fields of T4/T5 neurons. We fitted the parameters for the three axes of motion for both translation (Tx,Ty,Tz) and rotation (Rx,Ry,Rz) (**Fig. 4B**). Although the population data did not fully cover the visual field of one eye (**fig. S5A**), optic flow fields were well-matched filters for each of the six T4/T5 subtypes (**Fig. 4C** and **fig. S5B**). We compared performance including models where the fly only turned (rotational optic flow) or moved straight (translational optic flow). Across the six subtypes, only the model combining rotations and translations outperformed the null model consisting of a uniform vector field, with larger performance in layers A.I, B.II, C and layer D (**Fig. 4D** and **fig. S4C**). Thus, T4/T5 subtypes are tuned to optic flow generated by mixtures of translational and rotational motion. For each T4/T5 subtype, a single component tended to dominate the translational axis of motion, whereas the rotational axis was more distributed across the three components (**Fig. 4C** and **fig. S5A**). Together, our data show that local direction-selective T4/T5 neurons display a population code for different types of self-motion of the fly. Populations of T4/T5 cells are tuned to optic flow patterns, similar to their vertebrate counterparts(7), but representing six instead of four types of self-motion.

## Discussion

In this study, we have demonstrated that the local motion detectors T4 and T5 are divided into six subtypes that encode specific optic flow pattern. T4/T5 appear to implement an optimal population code for global motion patterns containing information about translational and rotational self-motion of the fly.

Direction-selective T4/T5 neurons in *Drosophila* have been described to encode four cardinal directions of motion(8, 9). Population T4/T5 recordings now reveal average tuning to diagonal rather than cardinal motion directions, such that six subtypes of T4/T5 neurons exist. Only a global analysis of directional tuning reveals these six subtypes, but tuning to diagonal motion has been observed in electrophysiological recordings of an individual T4 neuron(30), and in optical recordings of T4/T5(9, 31). T4/T5 neurons compute direction-selective signals across neighboring columns within the eye(32, 33). Thus, motion can simply be computed along the internal organization of the fly eye, and the hexagonal arrangement of the eye does not need to be transformed into a cardinal coordinate system.

Individual directional preference of a T4/T5 neuron correlates with its dendrite orientation, which manifests during development(34–36). Interestingly, developmental dendrite orientation for subtypes A and B reveal two peaks of diagonal rather than orthogonal orientation of dendrites(36), consistent with their distribution of direction selectivity. Our data show that each T4/T5 subtype retinotopically covers overlapping regions in visual space. A comprehensive analysis of T4/T5 dendrite anatomy across the visual system will be needed to clarify how adult dendrite orientation is distributed across the visual system to represent the six subtypes. Furthermore, single-cell transcriptomics has assigned developing T4/T5 cells to distinct clusters based on their genetic profiles(36–39), but genes involved in dendrite development or the differentiation are expressed in narrow time windows(39). Interestingly, one recent study identified a genetic subpopulation of T4 neurons, restricted to lobula plate layers A and B(37). While it remains to be determined whether this corresponds to the functional layer A/B subtypes, genetic access will help to better understand the development and anatomy of the individual subtypes.

While downstream of T4/T5, wide-field LPTCs are thought to encode self-motion(13, 17), our data show that the population of T4/T5 cells already encodes optic flow generated by a combination of rotational and translational self-motion of the fly. Within an optic flow field, single T4/T5 tuning changes along the retinotopic map. This could be inherited by the spatial distribution of ommatidia along the optical axis which varies with the curvature of the eye(40, 41). T4/T5 can then pass this information to downstream LPTCs, which do to not need to transform cardinal motion information into complex flow fields. Further internal dendritic processing, such as suppression of adjacent local motion signals, electrical coupling between LPTCs (17) and feedforward inhibition from lobula plate intrinsic neurons (23) will support the computation of diverse optic flow fields(22, 23, 26).

The fly eye and the vertebrate retina both show differences between local and global directional tuning(7, 42), and similarly compute visual signals generated by self-motion at the population level(7). A population code for optic flow generated by self-motion might therefore be canonical and evolved convergently during evolution. Such functionally driven convergence of neuronal circuit organization argues for an optimal design of encoding self-motion. However, mice and flies differ in the number and directions of optic flow encoded by local direction-selective cells. Flying animals might encode more motion axes than walking animals to match the higher degrees of freedom encountered during flight. This difference might therefore highlight adaptation to visuoecological niches of flying and walking animals. We are just starting to understand how a population code in visual systems matches the statistics of the visual environment(6, 7, 43–45), or animal behavior. Thus, this work is an important step towards understanding how anatomy, ethological constraints, and neuronal function are ultimately linked.

## Acknowledgements

We thank Carlotta Martelli, Yvette Fisher, Axel Methner, and members of the Silies lab for comments on the manuscript, and Christof Rickert for help with image analysis. This project has received funding from the European Research Council (ERC) under the European Union’s Horizon 2020 research and innovation program (grant agreement No 716512).

## Author contributions

MH and MS conceived the experiments. MH performed experiments and analyzed data. BG helped with data analysis. GRT modeled the flow fields. MH and MS wrote the paper.

## Materials and Methods

### *Drosophila* strains and fly husbandry

*Drosophila melanogaster* were raised on molasses-based food at 25°C and 55% humidity in a 12:12 hr light-dark cycle. For all imaging experiments female flies of the genotype *w*^*+*^; *R59E08-LexA*^*attP40*^, *lexAop-GCaMP6f-p10*^*su(Hw)attp5*^*/ R59E08-LexA*^*attP40*^, *lexAop-GCaMP6f-p10*^*su(Hw)attp5*^ were recorded 3-5 days after eclosion at room temperature (RT, 20°C). *R59E08-LexA*^*attP40*^ and *lexAop-GCaMP6f-p10*^*su(Hw)attp5*^ were obtained from the Bloomington Drosophila Stock Center (BDSC #52832 and #44277), recombined, and crossed into a *w*^*+*^ background.

### *In vivo* two-photon calcium imaging

#### Fly preparation, experimental setup and data acquisition

Prior to two-photon imaging, flies were anesthetized on ice and fit into a small hole in stainless-steel foil, located in a custom-made holder. The head was tilted approximately 30° to expose the back of the head. To fix the head of the fly, a small drop of UV-sensitive glue (Bondic) was used on the left side of the brain and the thorax. The cuticle on the right eye, fat bodies and tracheae were removed using breakable razor blades and forceps. To ensure constant nutrients and calcium supply flies were perfused with a carboxygenated saline containing 103 mM NaCl, 3 mM KCl, 5 mM TES, 1mM NaH2PO4, 4 mM MgCl2 1.5 mM CaCl2, 10mM trehalose, 10mM glucose, 7mM sucrose, and 26mM NaHCO3 (pH∼7.3). To record calcium activity, a two-photon microscope (Bruker Investigator, Bruker, Madison, WI, USA), equipped with a 25x/1.1 objective (Nikon, Minato, Japan) was used. For excitation of GCaMP6f, the excitation laser (Spectraphysics Insight DS+) was tuned to a wavelength of 920nm with *<*20mW of laser power measured at the objective. Emitted light was filtered through an SP680 short pass filter, a 560 lpxr dichroic filter and a 525/70 emission filter and detected by PMTs set to a gain of 855V. Imaging frames were acquired at a frame rate of ∼15-20 Hz and 4-7 optical zoom using PrairieView software. Each fly was recorded in at least three to five different focal planes (z-depth). We determined z-depth position relative to cell bodies and started the first recording at a z-depth of 30μm from there. Planes were then imaged every 15μm from there (Error! Reference source not found.**b**).

#### Visual stimulation

Visual stimuli were presented on an 8 cm x 8 cm rear projection screen in front of the fly covering a visual angle of 60° in azimuth and elevation. To cover a larger part of the horizontal visual field of 150° we rotated the fly with respect to the screen two times by 45° (Extended Data Fig. 1a). Stimuli were filtered through a 482/18 bandpass filter (Semrock) and ND1.0 neutral density filter (Thorlabs) and projected using a LightCrafter 4500 DLP (Texas Instruments, Texas, USA) with a frame rate of 100 Hz and synchronized with the recording of the microscope as described previously(46). All visual stimuli were generated using custom-written software using C++ and OpenGL.

#### Moving OFF and ON edges

Full-contrast dark or bright edges moving with a velocity of 20°/s across the full screen to four or eight different directions. Each stimulus direction was presented at least twice in pseudo-random order. The four-direction stimulus was merely used for the subsequent identification of T4 and T5 axon terminals.

### Data analysis

#### Preprocessing

All data analysis was performed using MATLAB R2017a (The MathWorks Inc, Natrick, MA) or Python 2.7. Motion artifacts were corrected using Sequential Image Alignment SIMA, applying an extended Hidden Markov Model(47).

#### Automated ROI selection

For the extraction of single T4 or T5 axon terminals we made use of their contrast- and direction-selective responses to ON and OFF edges moving into four directions. First, the aligned images were averaged across time and the average image intensity was Gaussian filtered (s=1.5) and then threshold-selected by Otsu’s method(48) to find foreground pixels suitable for further analysis. After averaging responses across stimulus repetitions, we selected pixels that showed a peak response larger than the average response plus two times the standard deviation of the full trace. These pixels were grouped based on their contrast preference (ON or OFF pixels) and further assigned to four categories based on their anatomical location within the lobula plate (layers A, B, C, or D). We further calculated a direction-selectivity index (DSI) and contrast selectivity index (CSI) for each pixel as follows:

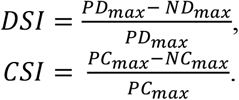

where *PD*_*max*_ and *ND*_*max*_ denote the maximal response into the preferred direction (PD) and null direction (ND) and *PC*_*max*_ and *NC*_*max*_ denote the maximum responses for the preferred contrast (PC) and the non-preferred or inverse contrast (NC). We excluded all pixels that did not exceed the CSI threshold of 0.2 to obtain clean T4 or T5 responses. For the final clustering we used the quantified DSI and CSI parameter and the timing of the response to the PD. Based on these parameters the Euclidean distance between each pair of pixels was calculated and average-linkage agglomerative hierarchical clustering was performed. We further evaluated the optimal distance threshold that yielded most clusters of the appropriate size between 1 and 2.5 μm^2^. All resulting clusters that fell outside this range were excluded from further analysis. Custer locations were saved and matched with subsequent recordings of the same cells to other stimulus types.

#### Moving OFF and ON stripes

For dF/F calculation, baseline responses to ∼ 0.5s gray epoch were used. To quantify direction selectivity (DS) of single cells, responses were trial averaged and the peak response to the eight different directions of either increment or decrement bars was extracted for T4 and T5 cells respectively. We further quantified the tuning of single cells by computing vector spaces as follows(27):

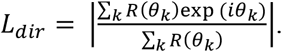

where *R*(*θ*_*k*_) is the response to angle *θ*_*k*_. The direction of the vector *L*_*dir*,_ denotes the tuning angle of the cell and the normalized length of the vector is related to the circular variance and thus represents the selectivity of the cell.

#### Receptive-field center extraction

To extract receptive-field centers, we used a back-propagation algorithm to map the receptive fields of T4 and T5 cells and to locate the center of the receptive fields(49). First, we imaged neural responses to eight different directions and created two-dimensional images from one-dimensional response traces. Neural latency and indicator dynamics introduce delays that will decrease the precision of receptive-field position estimation. To account for this delay, we measured the spatial difference of the response peaks between a static and a moving stimulus. We found an average of 9.6° delay for both T4 and T5 cells and shifted the traces for 9.6° before calculating the receptive-field map in our back-propagation algorithm (Error! Reference source not found.**a, b**). These were rotated according to their corresponding direction and averaged to obtain a receptive-field map. To find the center of the receptive field, we fitted a two-dimensional gaussian and took its peak coordinate.

#### Z-stack generation

Images representing the location of single ROIs color coded by their directional preference were generated in Matlab. Images containing data from different z-depth layers within the same fly were then further processed in Illustrator to create pseudo z-stacks. For this, ROIs from the same lobula plate layer were first compiled in a 3D structure and ROIs from different z-depth layers were stacked to better represent the third dimension of the lobula plate.

### Statistics

All statistics were done in Matlab using Circular Statistics Toolbox(50).

#### SNOB analysis

To extract underlying classes from the population of neurons found in layers A and B, we converted data of each population to be linear in the range of directions that most neurons where selective to, resulting in a scale from - π to π for layer A, C and D and a scale from 0 to 2π for the data from layer B. We used the finite mixture model SNOB(51) to predict the number of underlying Gaussians using minimum message length criterion. We further used the statistical prediction from the model to assign individual neurons to each of the underlying classes by choosing the class with the highest probability of the neuron’s tuning preference (Error! Reference source not found.**c**,**d**).

### Model

We fitted an optic flow field elicited from self-motion on the field of view at a constant distance from the observer, i.e., a spherical surface. Two coordinates describe the viewing direction: the azimuth *θ* and the elevation *ϕ* angles.

The self-motion flow-field vectors 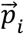 at each viewing location 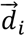 on the unit sphere were specified by the translation and rotation vectors, 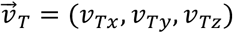 and 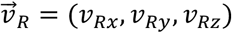, respectively (28):

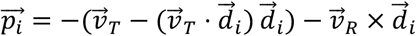

The flow-field vectors were then represented in spherical coordinates 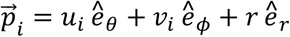 to extract a vector tangential to the spherical surface 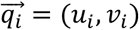 that could be matched to the direction-selectivity vectors from T4/T5 data. The uniform flow-field tangent to the spherical surface was specified by a single vector 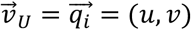 at every viewing position.

The comparison of data to model was done using the following *loss* function

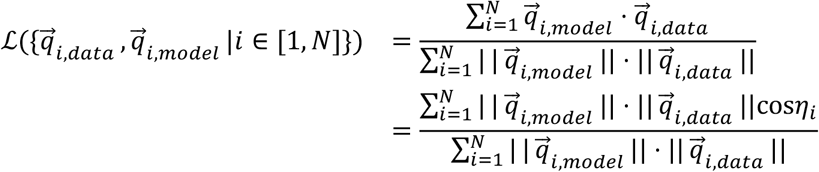

where *η*_*i*_ is the angle between the model and the data flow vectors at the location 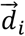, for all *N* vectors in the dataset, and ∥ · ∥ indicates the magnitude of the vector. When all vectors match in both magnitude and direction this quantity is 1 and when all vectors match in magnitude but are in opposite directions this quantity is -1. To optimize for the vectors 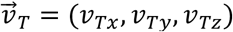 and 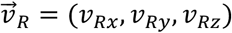, and 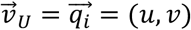 that maximize *ℒ*, the MATLAB function fmincon was used. The positive of the loss function is the linear projection (LP), shown in (**Fig. 4c**,**d**) and (**Error! Reference source not found.a**,**c**).

Four model variations were considered: fitting both 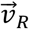 an 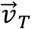; fitting 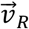 with 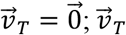; with 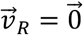; and fitting the uniform model 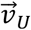. For all cases the model was constrained to vectors of unit magnitude, to focus on the direction rather than the speed of self-motion, and because the T4/T5 vectors (direction-selectivity index) had magnitudes between zero and one.

The data was fitted using ten-fold cross-validation (CV), dividing the data into ten random subsets. In each fold, nine subsets were used for training and the remaining subset was used for testing the model fit. For each CV fold, the same training data was fit ten times starting from ten different random conditions, and the best fit was stored and used to calculate the performance on the test set. The same training and testing data were used for all models, resulting in repeated measures of the test performance across models. Statistical testing was done on the ten test-performance values obtained per model. A one-tailed nonparametric Wilcoxon signed-rank test was used to determine whether the performance of each of the self-motion models was higher than the performance of the uniform model, for tests pooling all subtypes (**Fig. 4d**) and tests of individual subtypes (**Error! Reference source not found.c**).A signed-rank test accounted for repeated measures, and a Bonferroni correction was applied to account for multiple testing (*p <* 0.05/3).

